# Optimization of an Experimental Vaccine to Prevent *Escherichia coli* Urinary Tract Infection

**DOI:** 10.1101/2020.03.10.986729

**Authors:** Valerie S. Forsyth, Stephanie D. Himpsl, Sara N. Smith, Christina A. Sarkissian, Laura A. Mike, Jolie A. Stocki, Anna Sintsova, Christopher J. Alteri, Harry L.T. Mobley

## Abstract

Urinary tract infections (UTI) affect half of all women at least once during their lifetime. The rise in extended-spectrum beta-lactamase-producing strains and potential for carbapenem resistance within uropathogenic *Escherichia coli* (UPEC), the most common causative agent of UTIs, creates an urgent need for vaccine development. Intranasal immunization of mice with UPEC outer membrane iron receptors, FyuA, Hma, IreA, or IutA, conjugated to cholera toxin, provides protection in the bladder or kidneys when challenged with UPEC CFT073 or 536. Based on these data, we sought to optimize the vaccination route (intramuscular, intranasal, or subcutaneous) in combination with adjuvants suitable for human use including alum, monophosphoryl lipid A (MPLA), unmethylated CpG synthetic oligodeoxynucleotides (CpG), polyinosinic:polycytodylic acid (polyIC), and mutated heat-labile *E. coli* enterotoxin (dmLT). Mice intranasally vaccinated with dmLT-IutA or dmLT-Hma displayed a significant reduction in bladder colonization (86-fold and 32-fold, respectively) with 40–42% of mice having no detectable colony forming units (CFU). Intranasal vaccination of mice with CpG-IutA and polyIC-IutA significantly reduced kidney colonization (131-fold) and urine CFU (22-fold), respectively. dmLT generated the most consistently robust antibody response in intranasally immunized mice, while MPLA and alum produced greater concentrations of antigen-specific serum IgG with intramuscular immunization. Based on these results, we conclude that intranasal administration of Hma or IutA formulated with dmLT adjuvant provides the greatest protection from UPEC UTI. This study advances our progress toward a vaccine against uncomplicated UTI, which will significantly improve the quality of life for women burdened by recurrent UTI and enable better antibiotic stewardship.

**Importance:** Urinary tract infections (UTI) are among the most common bacterial infection in humans, affecting half of all women at least once during their lifetimes. The rise in antibiotic resistance and health care costs emphasizes the need to develop a vaccine against the most common UTI pathogen, *Escherichia coli.* Vaccinating mice intranasally with a detoxified heat-labile enterotoxin and two surface exposed receptors, Hma or IutA, significantly reduced bacterial burden in the bladder. This work highlights progress in the development of a UTI vaccine formulated with adjuvants suitable for human use and antigens that encode outer membrane iron receptors required for infection in the iron-limited urinary tract.

## Introduction

Urinary tract infections (UTI), the second most common human infection after respiratory infections, result in an annual cost of $ 3.5 billion (1, 2). Uropathogenic *Escherichia coli* (UPEC) is the most prevalent causative agent of uncomplicated UTI, rates of antibiotic resistance in pathogenic isolates are increasing, and multidrug resistant strains (*E. coli* ST131) are emerging (1–3). Despite innate immune defenses in the bladder that include micturition, a mucin layer, constitutively expressed secretory immunoglobulin A, cationic antimicrobial peptides, Tamm-Horsfall protein, lactoferrin, and lipocalin-2 (4), half of all women will experience a UTI in their lifetime with 1 in 40 women experiencing recurrent infections (5). Patients with acute or recurrent UTI have significantly decreased levels of total secretory IgA in the urine as compared to healthy individuals with no history of UTI (6, 7). This indicates the potential for decreased severity and duration of infection if microbe-specific antibody levels can be increased with a vaccine. Because 90% of symptomatic UTI are uncomplicated infections, an ideal vaccine will target factors critical for establishment of bladder colonization (3) Five FDA-approved vaccines provide mucosal protection against other pathogens including poliovirus, rotavirus, influenza virus, *Salmonella enterica* serovar Typhi, and *Vibrio cholerae* (8–11). These efficacious mucosal vaccines that protect against other enteric viruses and bacteria bolster the hypothesis that a vaccine effective against uropathogens is attainable.

During the last 20 years there have been noteworthy advancements toward the development of a UTI vaccine, yet no licensed UTI vaccines are available for use in the U.S. Published studies have investigated the efficacy of vaccines containing O antigen (12), fimbrial subunits (13, 14), α-hemolysin (15), siderophores (16), and a variety of outer membrane siderophore receptors in animal models of UTI (17–20). Human clinical trials have been performed on three vaccines, Uro-Vaxom, SolcoUrovac, and ExPEC4V. Uro-Vaxom, comprised of 18 *E. coli* uropathogen extracts and administered as a daily oral tablet, is approved in Germany and Switzerland for the prevention of recurrent cystitis (21). SolcoUrovac, currently marketed as StroVac, contains heat-killed uropathogenic bacteria including *E. coli, Proteus vulgaris, Klebsiella pneumoniae, Morganella morganii*, and *Enterococcus faecalis* and is approved for human use in Europe (3, 22). ExPEC4V consists of four conjugated O-antigens O1A, O2, O6A, and O25B common to *E. coli* strains known to cause UTI (23). In a study comparing the efficacy of these three vaccines in adults with recurrent UTIs, Uro-Vaxom had the greatest reduction in rate of UTI recurrence while ExPEC4V did not appear to reduce UTI recurrence (24). Nonetheless, the daily regimen and toxic side effects have limited widespread use of Uro-Vaxom (25).

Here we describe our efforts to develop a vaccine against uncomplicated UTIs using antigens previously identified and validated as vaccine candidates by intranasal immunization in a murine UTI model when conjugated to the adjuvant cholera toxin (26–28). We previously performed an extensive multi-omics approach to identify genes and their proteins that: 1) are localized to the bacterial cell surface (29); 2) are expressed during growth in human urine (30), murine infection (31), and human infection (32, 33); 3) possess immunoreactive properties (34); and 4) are more prevalent in UPEC isolates than commensal isolates (35, 36). A total of four β-barrel outer membrane receptors required for iron sequestration met all of these criteria including heme receptor Hma, aerobactin receptor IutA, yersiniabactin receptor FyuA, and putative siderophore receptor IreA. Effective iron acquisition from the iron-limited environment of the urinary tract is required for full virulence of UPEC (37–39). In addition to their iron scavenging function, IreA functions as an adhesin that is important for colonization of the bladder (38) and FyuA plays a role in biofilm formation in human urine (40). Intranasal immunization with Hma, IreA, IutA, or FyuA, conjugated to cholera toxin, significantly reduced in bacterial burden in the bladder, kidneys or both 48 hours following transurethral challenge with UPEC (27, 28). While cholera toxin is an effective immune stimulant in mice, it is not suitable for human use, due to development of profuse diarrhea with oral doses as low as 5 µg (41). Because of this drawback, we sought to optimize this UTI vaccine by incorporating adjuvants approved for use in humans or used in vaccine clinical trials.

The precise immune response required for protection against UTI is not well-defined. Therefore, we selected a panel of adjuvants known to elicit an array of adaptive immune responses, with the aim being to identify an adjuvant that is well-suited for protecting against UTI and safe for use in humans (3, 42). The five adjuvants tested were alum (43, 44), monophosphoryl lipid A (MPLA) (45–47), unmethylated CpG synthetic oligodeoxynucleotides (CpG) (48–50), polyinosinic:polycytodylic acid (polyIC) (51), and double mutant (R192G/L211A) heat-labile *E. coli* enterotoxin (dmLT) (52). Alum is licensed for use in twenty-two vaccines available in the U.S. and is reported to activate dendritic cells via multiple mechanisms, thus promoting antigen uptake and release of IL-1β and IL-18 (43, 53). MPLA is derived from *Salmonella minnesota* R595 lipopolysaccharide, activates cellular immunity through the TLR4 signaling pathway, is approved for human use in Europe, and is a component of vaccines for hepatitis B and papilloma viruses (51, 54). CpG activates TLR9 signaling in B cells and dendritic cells, increases mucosal immune responses, and is licensed for use in a hepatitis B vaccine (46, 47, 55). Both dmLT and CpG are presumed to function by activating innate signaling and stimulation of mucosal dendritic cells, which activate the adaptive immune response, particularly Th17 cells by dmLT (52, 56–58). polyIC is a synthetically produced double stranded RNA, analogous to viral RNA, that induces a robust type I interferon response resulting in activation of cellular immunity, and is in late stage clinical development (51).

In an effort to develop a vaccine protective against uncomplicated UTI in humans, we tested five adjuvants (dmLT, CpG, polyIC, MPLA, and alum) with four antigens (Hma, IreA, IutA, or FyuA) for efficacy in mice. Because immunization route can affect the immune response, we examined three routes of immunization: intranasal, intramuscular, and subcutaneous with multiple antigen-adjuvant combinations. Hma and IreA, which have been shown to significantly reducepreviously demonstrated to provide the most robust reduction of bacterial burden in the kidneys and bladder (28), respectively, were initially examined under all conditions, then the most promising combinations of route and adjuvant were further evaluated for efficacy with the remaining two antigens FyuA and IutA. Here we report that intranasal immunization with dmLT-Hma and dmLT-IutA induces antigen-specific antibody production and provides robust protection in immunized mice following transurethral challenge with UPEC.

## Results

### Immunizing via the intranasal route provides the most protection against UTI

We previously established that intranasal immunization followed by two weekly boosts with outer membrane iron receptors Hma, IreA, IutA or FyuA conjugated to cholera toxin significantly reduced bacterial burden in the bladder, kidneys or both, 48 hours following transurethral challenge with UPEC (27, 28). We later determined that conjugation of antigen to adjuvant was not required for protection (unpublished results). Based on these data, we began systematic optimization of the vaccine to maximize efficacy using alternative adjuvants admixed with antigen. The optimized vaccine route, adjuvant and antigen were evaluated based on three criteria: reduction of bacterial CFU in the sample sites, increased number of mice without detectable bacterial counts, and production of antigen-specific antibody. Genes encoding all protein antigens were codon-optimized, proteins were purified from inclusion bodies and certified LPS-free.

The immunization route can markedly affect the efficacy of vaccines, and some routes are linked to greater patient compliance especially when immunization requires multiple boosts (59, 60). In an effort to determine the optimal route, we systematically immunized mice with one of five adjuvants either intranasally, intramuscularly, or subcutaneously in combination with either Hma or IreA, which have been shown to significantly reduce bacterial burden in the kidneys and bladder, respectively (28). Promising combinations of route and adjuvant were further evaluated for efficacy by imunizing with the remaining two antigens FyuA and IutA. Intramuscular and subcutaneous routes were chosen to expand upon early success with intranasal immunization because they are commonly used deliver vaccines in humans. Mice were immunized as previously described (28). In total, 35 immunization trials utilizing 1,060 mice were performed.

We found that intranasal immunization tended to reduce median bacterial burden at least two-fold in the urine for all antigen-adjuvant combinations tested (Table 1). When immunized intranasally, two-fold reduction in median bacterial burden in the bladder occurred for 58% (7/12) of combinations tested and in the kidneys for 33% (4/12) of combinations tested (Table 1). Significant differences are noted in bold in Table 1. Specifically, intranasal immunization with polyIC-IutA (*P*=0.059 urine), dmLT-Hma (*P*=0.024 bladder), dmLT-IutA (*P*=0.018 bladder) and CpG-IutA (*P*=0.046 kidneys) significantly reduced bacterial burden (Figure 1B, E, F, J, Table 1). Three adjuvant-antigen combinations tended to reduce colony forming units (CFU) more than two-fold in all sites of infection following intranasal immunization: dmLT-IutA, CpG-Hma, and CpG-IreA (Table 1). The bimodal distribution of bladder colonization observed with protective combinations (dmLT-Hma, dmLT-IutA) is typical of this model (19, 20).

**Table 1.** Median fold change in CFU in the urine, bladder and kidneys of immunized mice.

| Route | Adjuvant | Antigen | Urine |  |  |  | Bladder |  |  |  | Kidneys |  |  |  |
| --- | --- | --- | --- | --- | --- | --- | --- | --- | --- | --- | --- | --- | --- | --- |
|  |  |  | Median Adj Only<sup>a</sup> | Median Adj + Antigen<sup>b</sup> | Fold Change<sup>c</sup> | P value<sup>d</sup> | Median Adj Only | Median Adj + Antigen | Fold Change | P value | Median Adj Only | Median Adj + Antigen | Fold Change | P value |
| IM<sup>e</sup> | alum | Hma | 1840 | 105300 | 0.02 | **0.0257** | 100 | 8670 | 0.01 | 0.2628 | 10940 | 6860 | 1.59 | 0.9861 |
| IM | alum | IreA | 1840 | 7285 | 0.25 | 0.0664 | 100 | 2950 | 0.03 | 0.9709 | 10940 | 10045 | 1.09 | 0.5761 |
| IM | polyIC<sup>f</sup> | Hma | 23100 | 734500 | 0.03 | 0.4973 | 6735 | 37350 | 0.18 | **0.0522** | 9650 | 3975 | **2.43** | 0.2834 |
| IM | polyIC | IreA | 23100 | 106000 | 0.22 | 0.6421 | 6735 | 13550 | 0.50 | 0.5326 | 9650 | 750 | **12.87** | 0.1808 |
| IM | MPLA<sup>g</sup> | Hma | 105350 | 562500 | 0.19 | 0.4813 | 100 | 4810 | 0.02 | 0.3285 | 3455 | 13620 | 0.25 | 0.3888 |
| IM | MPLA | IreA | 105350 | 17100 | **6.16** | 0.2466 | 100 | 1690 | 0.06 | 0.7799 | 3455 | 470 | **7.35** | 0.8994 |
| IM | dmLT<sup>h</sup> | Hma | 218500 | 101100 | **2.16** | 0.3440 | 28200 | 92650 | 0.30 | 0.4311 | 40400 | 6650 | **6.08** | 0.2400 |
| IM | dmLT | IreA | 218500 | 36500 | **5.99** | 0.1127 | 28200 | 14500 | 1.94 | 0.7275 | 40400 | 11000 | **3.67** | 0.7252 |
| IN<sup>i</sup> | polyIC | Hma | 10950 | 3500 | **3.13** | 0.2409 | 14600 | 6355 | **2.30** | 0.1009 | 1409 | 1810 | 0.78 | >0.9999 |
| IN | polyIC | IreA | 10950 | 4845 | **2.26** | 0.4122 | 14600 | 3330 | **4.38** | 0.1457 | 1409 | 5125 | 0.27 | 0.7937 |
| IN | polyIC | FyuA | 284000 | 54100 | **5.25** | 0.3097 | 5500 | 11800 | 0.47 | 0.5941 | 1180 | 927 | 1.27 | >0.9999 |
| IN | polyIC | IutA | 1440000 | 66700 | **21.59** | **0.0599** | 50900 | 11800 | **4.31** | 0.1469 | 640 | 1600 | 0.40 | 0.5804 |
| IN | dmLT | Hma | 2790 | 100 | **27.90** | 0.0693 | 3205 | 100 | **32.05** | **0.0240** | 407 | 542 | 0.75 | 0.6033 |
| IN | dmLT | IreA | 2790 | 105 | **26.57** | 0.3674 | 3205 | 3170 | 1.01 | 0.9020 | 407 | 1340 | 0.30 | 0.7087 |
| IN | dmLT | FyuA | 210450 | 5605 | **37.55** | 0.0825 | 2955 | 2835 | 1.04 | 0.4233 | 387 | 3159 | 0.12 | **0.0476** |
| IN | dmLT | IutA | 16400 | 619 | **26.49** | 0.1866 | 8610 | 100 | **86.10** | **0.0181** | 4950 | 1130 | **4.38** | 0.2536 |
| IN | CpG<sup>j</sup> | Hma | 97050 | 9285 | **10.45** | 0.4332 | 19350 | 5755 | **3.36** | 0.1321 | 13500 | 1885 | **7.16** | 0.1419 |
| IN | CpG | IreA | 97050 | 8215 | **11.81** | 0.4944 | 19350 | 3035 | **6.38** | 0.0608 | 13500 | 6490 | **2.08** | 0.2991 |
| IN | CpG | FyuA | 7910 | 961 | **8.23** | 0.6403 | 3130 | 5625 | 0.56 | 0.8550 | 312 | 773 | 0.40 | 0.8330 |
| IN | CpG | IutA | 44530 | 1240 | **35.91** | 0.1959 | 5255 | 8570 | 0.61 | 0.7791 | 21100 | 161 | **131.06** | **0.0462** |
| SQ<sup>k</sup> | alum | Hma | 2218 | 100 | **22.18** | 0.1246 | 3240 | 8205 | 0.39 | 0.7940 | 198 | 100 | 1.98 | 0.2391 |
| SQ | alum | IreA | 2218 | 3165 | 0.70 | 0.7828 | 3240 | 4655 | 0.70 | 0.9271 | 198 | 2315 | 0.09 | 0.3376 |
| SQ | polyIC | Hma | 100 | 1705 | 0.06 | 0.7168 | 1505 | 14650 | 0.10 | **0.0052** | 100 | 100 | 1.00 | 0.4241 |
| SQ | polyIC | IreA | 100 | 1710 | 0.06 | 0.5473 | 1505 | 4410 | 0.34 | 0.2559 | 100 | 9615 | 0.01 | **0.0274** |
| SQ | dmLT | Hma | 119000 | 8955 | **13.29** | 0.3154 | 2750 | 2555 | 1.08 | 0.9478 | 4390 | 12440 | 0.35 | 0.4422 |
| SQ | dmLT | IreA | 119000 | 213500 | 0.56 | 0.9048 | 2750 | 11360 | 0.24 | 0.7122 | 4390 | 4990 | 0.88 | 0.9675 |
| SQ | MPLA | Hma | 31900 | 232600 | 0.14 | 0.7988 | 5240 | 13800 | 0.38 | 0.5935 | 41800 | 5615 | **7.44** | 0.2818 |
| SQ | MPLA | IreA | 31900 | 202000 | 0.16 | 0.4002 | 5240 | 14700 | 0.36 | 0.0810 | 41800 | 54650 | 0.76 | 0.2309 |

<sup>a</sup>Median CFU when immunized with the adjuvant alone.
<sup>b</sup>Median CFU when immunized with the adjuvant formulated with antigen.
<sup>c</sup>Fold change in median CFU when immunized with the adjuvant alone compared to mice immunized with the adjuvant formulated with antigen. Fold change greater than 2 are shown in bold.
<sup>d</sup>P value as determined by two tailed Mann-Whitney exact test. Significant differences are shown in bold.
<sup>e</sup>Intramuscular <sup>f</sup>Polyinosinic:polycytidylic acid <sup>g</sup>Monophosphoryl lipid A <sup>h</sup>Detoxified *E. coli* enterotoxin <sup>i</sup>Intranasal <sup>j</sup>Unmethylated CpG synthetic oligodeoxynucleotides <sup>k</sup>Subcutaneous

**Figure 1.**
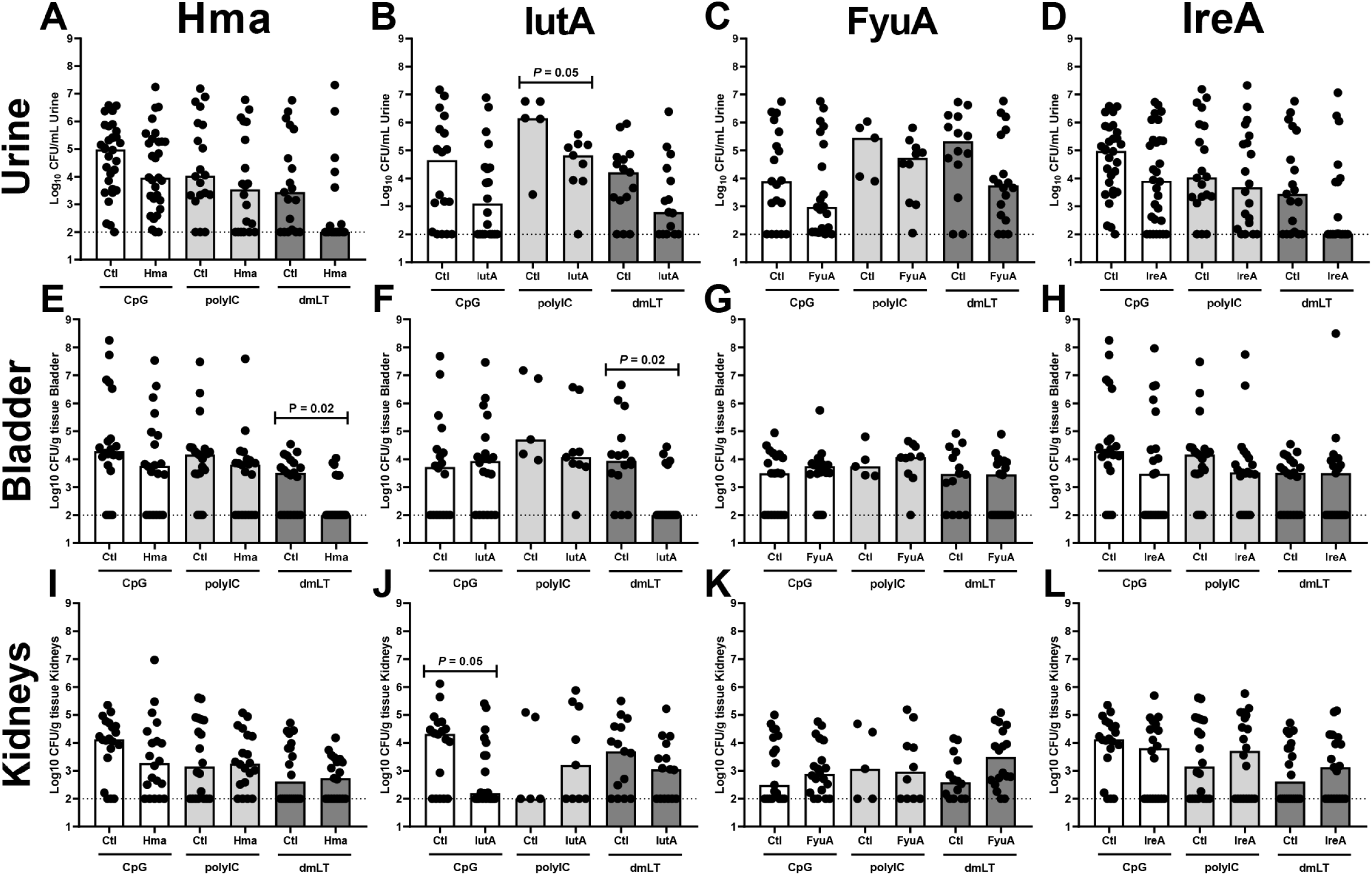
Comparison of adjuvants administered intranasally and formulated with the antigens Hma, IutA, FyuA, or IreA. Seven to eight-week old female CBA/J mice were immunized intranasally according to our immunization schedule with adjuvant alone (Ctl) or adjuvant formulated with 100 µg LPS free antigen, either Hma (A, E, I), IutA (B, F, J), FyuA (C, G, K), or IreA (D, H, L). Adjuvants tested included unmethylated CpG synthetic oligodeoxynucleotides (CpG), polyinosinic:polycytodylic acid (polyIC), or detoxified *E. coli* enterotoxin (dmLT). One week after the final boost, mice were challenged with 10^8^ colony forming units (CFU) of *E. coli* strain CFT073 (Hma, IreA, IutA) or 536 (FyuA) via transurethral inoculation. Forty-eight hours post inoculation, urine was collected, mice were sacrificed, bladder and kidneys homogenized, and aliquots plated on LB agar for enumeration of bacterial burden. Bars indicate the median CFU in the urine (A - D), bladder (E - H), and kidneys (I – L). Symbols represent individual mice. N = 5 - 30. Dashed line represents the limit of detection. *P* values were determined using a two-tailed Mann-Whitney test.

Immunization via the intramuscular and subcutaneous routes showed limited protection. Intramuscular immunization utilizing Hma as antigen tended to reduce median bacterial burden at least two-fold in 3 out of 12 trials (Table 1). Immunization with IreA intramuscularly did not reduce bacterial burden two-fold or greater in any trial. The subcutaneous route in combination with Hma tended to reduce median CFU at least two-fold in the urine when formulated with alum or dmLT, and in the kidneys when formulated with MPLA (Table 1). No reduction in bacterial burden was observed for any of the sites of infection when any of the IreA vaccine formulations were administered subcutaneously (Table 1).

Bacterial inocula for the thirty-five infection trials were confirmed to be at the desired dose with the median dose at 3.05 ± 0.11×10^9^ CFU/mL of strain CFT073 and 2.70 ± 0.20×10^9^ CFU/mL of strain 536 (Supplemental Figure 1). Strain CFT073 lacks a functional FyuA, therefore vaccine combinations containing this antigen utilized *E. coli* strain 536 for challenge. These data verify that variances in colonization level between the experimental trials were not due to differences in bacterial dose used to transurethrally inoculate mice (Supplemental Figure 1). When polyIC or CpG was administered intranasally in the absence of antigen, the median colonization level of CFT073 was not significantly different from that of unimmunized mice infected with the same dose of CFT073 (Supplemental Figure 2), indicating that polyIC or CpG alone do not alter colonization of CFT073. However, intranasal administration of dmLT as adjuvant in the absence of antigen significantly reduced the bacterial burden in the kidneys (*P*=0.02) (Supplemental Figure 2), consistent with previous findings that dmLT alone reduced colonization of multiple pathogens including *Haemophilis influenzae, Campylobacter jejuni*, and *Shigella flexneri* (52).

Taken together these results indicate that intranasal immunization tended to improve the protective response in the urine, bladder and kidneys at least two fold following bacterial challenge for 64% (23/36) of the adjuvant-antigen combinations, five of these were statistically significant, in comparison to intramuscular immunization 33% (8/24, *P* = 0.03) or subcutaneous immunization 13% (3/24, *P* < 0.005) (Table 1). No statistically significant CFU reductions were observed in any combination administered by the intramuscular or subcutaneous routes. Therefore, intranasal immunization is the optimal route for protection against transurethral challenge with *E. coli*.

### dmLT is the adjuvant that provides the most effective protection against colonization by *E. coli*

Having determined that intranasally administering different vaccine formulations provides the most consistent protection against UTI, we next set out to optimize the adjuvant. Previous studies have shown that immunizing with cholera toxin conjugated to antigens provides a robust immune response and reduces bacterial burden in the bladder and kidneys of CBA/J mice (27, 28), however, it is not suitable for human use (41). In an effort to develop a vaccine to prevent UTI, we tested our antigen candidates admixed with alternative adjuvants approved for human immunization: dmLT, CpG, polyIC, MPLA, and alum, for efficacy in mice. Intranasal immunization with dmLT, an adjuvant very similar in structure to cholera toxin (both are A_1_B_5_ toxins), significantly reduced the median bladder colonization (*P* = 0.02) when admixed with the antigen Hma compared to dmLT alone (Figure 1E). In addition, this vaccine combination significantly increased the number of mice without detectable bacteria in the bladder (35% dmLT only, 68% dmLT-Hma, *P* = 0.056), and urine (26% dmLT only, 61% dmLT-Hma, *P* = 0.05) indicating that more mice cleared the infection (below the limit of detection) 48 hours post-inoculation (Supplemental Table 1). Similar to Hma, immunizing with dmLT-IutA significantly reduced the median CFU/g bladder (*P* = 0.02) (Figure 1F) and significantly increased the number of protected mice (20% dmLT only, 67% dmLT-IutA, *P* = 0.03) (Supplemental Table 1). In addition, dmLT administered with FyuA and IutA tended to reduce urine bacterial load two-fold (Table 1).

Other adjuvant-antige combinations that resulted in significant reductions in bacterial burden when administered intranasally include polyIC-IutA (*P* = 0.05 urine) (Figure 1B) and CpG-IutA (*P* = 0.05 kidneys) (Figure 1J). The primary target of this vaccine is patients with uncomplicated UTI, therefore, reducing bladder colonization and inflammation (cystitis) is the desired outcome. Since dmLT-IutA and dmLT-Hma both reduced bacterial burdens in the bladder, dmLT was selected for future studies.

### IutA is the optimal antigen when co-administered with dmLT via the intranasal route

Previous studies with cholera toxin conjugated to antigens found that IreA and IutA provided significant protection from bacterial challenge in the bladder and Hma, FyuA, and IutA provided protection in the kidneys (27, 28). When administered via the intranasal route and formulated with dmLT, individually all four antigens trended toward reducing bacterial burdens in the urine (Figure 1A-D). In the bladder, CFU/g tissue was significantly reduced compared to adjuvant alone when Hma (32 fold-reduction, *P*=0.024) and IutA (86 fold-reduction, *P*=0.018) were co-administered with dmLT (Figure 1E, F). FyuA and IreA did not provide any protection compared to the adjuvant alone cohort (Table 1, Figure 1G, H). IutA was the only antigen that reduced colonization more than two-fold in the kidneys, although this reduction was not statistically significant (Table 1). According to these results, IutA is an effective antigen for reduction of bacterial burden when administered via the intranasal route in combination with dmLT in all sites of infection. In comparison Hma is an effective antigen in two sites of infection.

### Antigen-specific antibody response to immunization

In addition to evaluating the vaccine efficacy by observing changes in the bacterial burden after challenge, the humoral response was evaluated using an indirect ELISA. The amount of antigen-specific IgG in serum collected one week after the second boost was quantified. In all trials, independent of route, adjuvant or antigen, mice immunized with adjuvant alone had no measurable amounts of antigen-specific antibody (Figure 2, 3 Supplemental Figure 3). However, mice immunized with adjuvant admixed with antigen elicited a statistically significant increase in the concentration of antigen-specific IgG in the serum compared to controls immunized with adjuvant alone (*P* = 0.05) (Figure 2, 3 Supplemental Figure 3). This indicated that addition of adjuvant alone does not produce an antigen-specific immune response, and outer-membrane iron receptor preparations containing no LPS contamination are antigenic.

**Figure 2.**
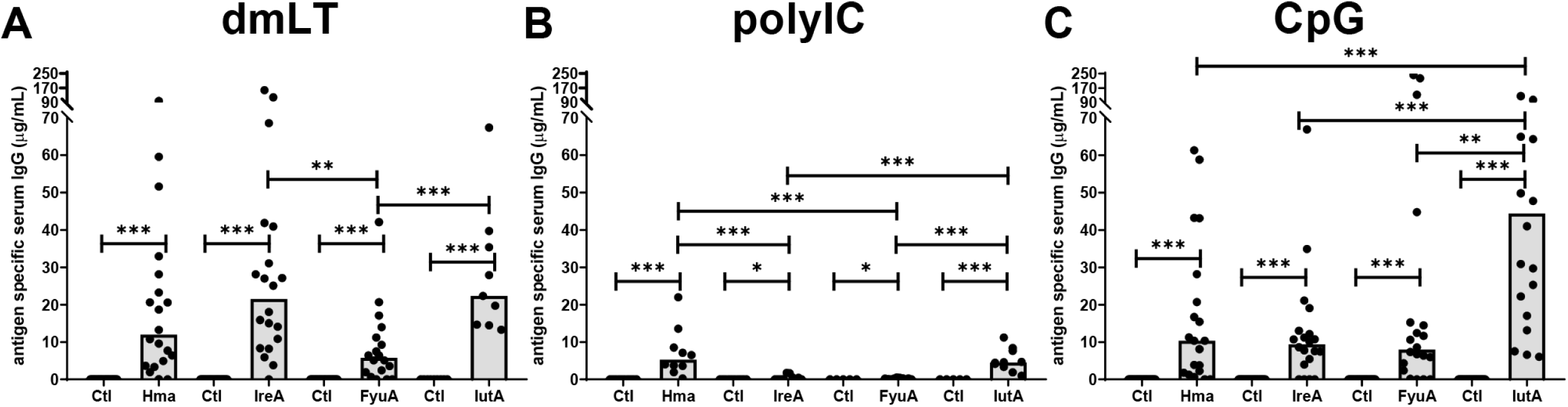
Intranasal vaccination with outer membrane iron receptors generates a robust antigen-specific serum IgG response. Antigen-specific IgG concentrations quantified by indirect ELISA in serum collected from female CBA/J mice one week after final boost. Mice were intranasally immunized with adjuvant alone (Ctl) or adjuvant formulated with 100 µg of purified, LPS free antigen (Hma, IreA, FyuA, IutA). Adjuvants utilized were (A) detoxified *E. coli* enterotoxin, dmLT, (B) polyinosinic:polycytodylic acid, polyIC, and (C) unmethylated CpG synthetic oligodeoxynucleotides, CpG. Each bar represents the median and each symbol represents an individual mouse. N = 5 – 20. *P* values were determined using a two-tailed Mann-Whitney test. *** *P* ≤0.001, ** *P* ≤0.01, * *P* ≤0.05.

**Figure 3.**
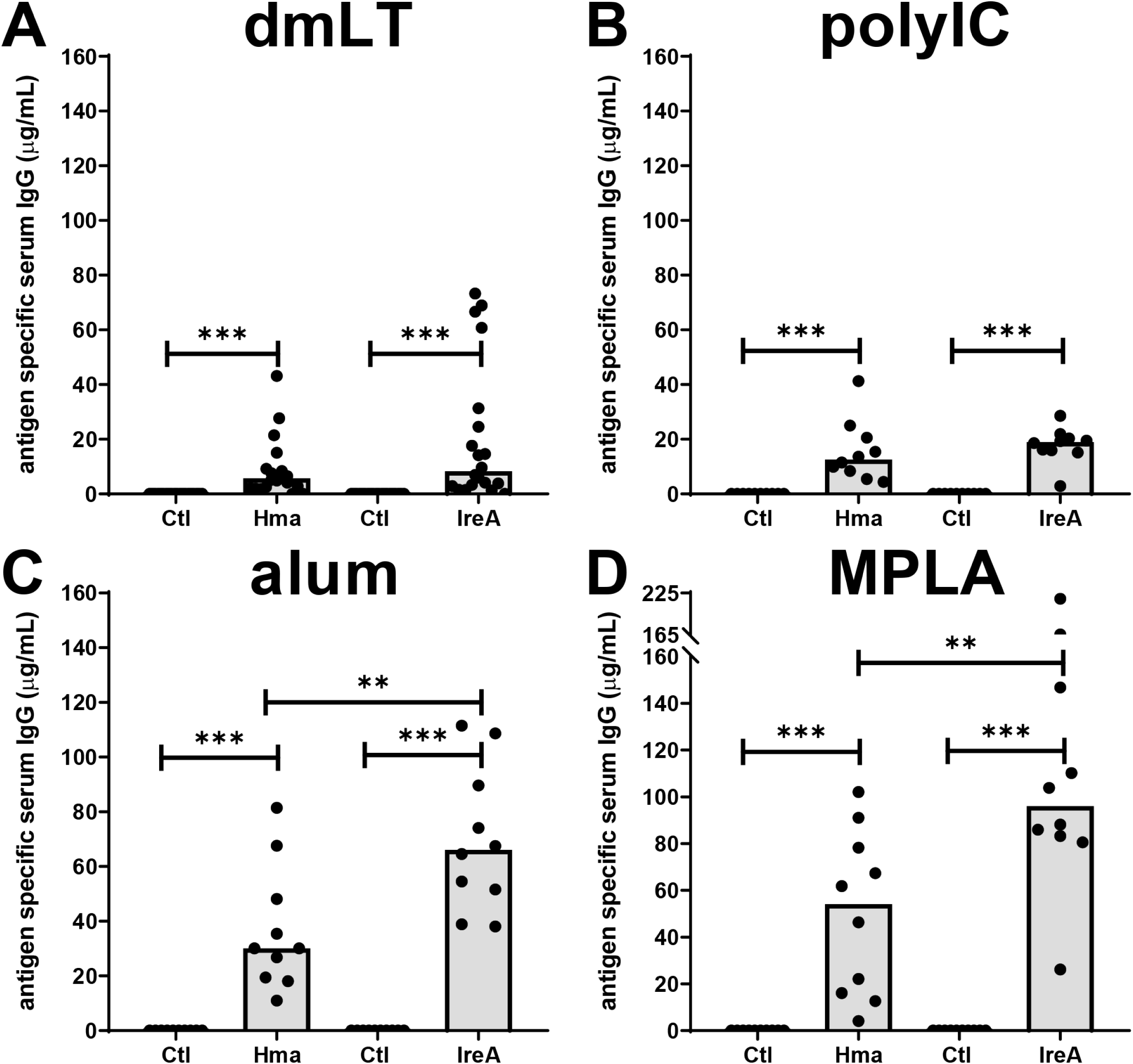
Intramuscular vaccination with outer membrane iron receptors generates a robust antigen-specific serum IgG response when alum or MPLA are used as adjuvants. Antigen-specific IgG concentrations quantified by indirect ELISA in serum collected from female CBA/J mice one week after final boost. Mice were intramuscularly immunized with adjuvant alone (Ctl) or adjuvant formulated with 100 µg of purified, LPS free antigen (Hma, IreA, FyuA, IutA). Adjuvants utilized were (A) detoxified *E. coli* enterotoxin, dmLT, (B) polyinosinic:polycytodylic acid, polyIC, (C) alum, and (D) monophosphoryl lipid A, MPLA. Note the change in scale in panel D. Each bar represents the median and each symbol represents an individual mouse. N = 10 – 20. *P* values were determined using a two-tailed Mann-Whitney test. *** *P* ≤0.001, ** *P* ≤0.01.

The immune responses generated between routes of immunization and adjuvants varied. Intranasal immunization with dmLT generated the most consistent and robust serum antibody response across all antigens, with median concentration of antigen-specific IgG 12.0 µg/mL Hma, 21.6 µg/mL IreA, 5.8 µg/mL FyuA, 22.4 µg/mL IutA (Figure 2A). When compared to vaccine formulations that significantly reduced bacterial burden in the bladder of challenged mice, there was no correlation between antigen-specific antibody production and protection for individual mice, indicating that the mechanism of protection may include cell-mediated immunity (Supplemental Figure 4). In comparison to immunizations with dmLT, polyIC in combination with all antigens produced a weak antigen-specific antibody response with an average concentration of 3.2 µg/mL serum IgG (Figure 2B). When all intranasal immunizations are compared, mice immunized intranasally with CpG-IutA produced the greatest median concentration of antigen-specific antibody of all intranasal immunizations (44.9 µg/mL), although the response was highly variable between individual mice (Figure 2C) and did not correlate with a reduction in bladder bacterial burden (Supplemental Figure 3D).

Although intramuscular and subcutaneous immunization were less effective at reducing the bacterial burden in the urine, bladder and kidneys of challenged mice, these routes were able to generate an antigen-specific IgG response in the serum (Figure 3, Supplemental Figure 3). Intramuscular immunizations with MPLA produced the highest median concentration of antigen-specific antibody for all vaccine trials (median Hma 54.1 µg/mL, IreA 96.0 µg/mL) (Figure 3D), with vaccine combinations that used alum as adjuvant producing similar responses (Figure 3C). Immunizations performed with dmLT as adjuvant had a very weak IgG response when administered intramuscularly (median Hma 5.7 µg/mL, IreA 8.3 µg/mL) (Figure 3A). Subcutaneous immunizations produced a similar antigen-specific IgG response independent of adjuvant, mean concentrations in pooled mouse sera ranging from 37.8 µg/mL to 97.3 µg/mL (Supplemental Figure 3). Most notably, for both the intramuscular and subcutaneous routes, immunization of mice with IreA increased the concentration of antigen-specific antibody above similar vaccine formulations where Hma was used as antigen (Figure 3, Supplemental Figure 3), indicating that IreA may be more antigenic despite having a highly similar tertiary structure.

## Discussion

Currently, there is no licensed vaccine in the U.S to prevent UTI by *E. coli*. In this current era of increasing antibiotic resistance, developing preventive measures against urinary tract pathogens, especially UPEC, is critical. Towards that goal, here we have systematically tested four antigens, previously validated for their protective efficacy against colonization of the bladder and kidneys during experimental murine UTI (27, 28) in combination with each of five adjuvants, compatible with eventual clinical trial design, delivered by each of three routes. Intranasal immunization was previously protective in mice, while intramuscular or subcutaneous routes are more commonly used in humans. Intranasal administration of dmLT formulated with Hma or IutA significantly reduced bladder CFU and significantly increased the number of mice without detectable CFU. Immunization with dmLT which, like cholera toxin, is an A_1_B_5_ toxin, consistently generated a robust IgG antibody response against the administered antigen. Optimization of the route, adjuvant and antigen is an important step in development of a UTI vaccine for human clinical trials.

Induction of mucosal immune responses is most efficiently stimulated by delivery of a vaccine at a mucosal surface (61) and may be due to the common mucosal immune system that links tissues of the lung, gastrointestinal tract, urogenital tract, and nasopharynx (62). In mice, monkeys, and humans, vaccines administered intranasally generate mucosal immune responses in the female genital tract (63–66). This is attributed to the expression of redundant B and T cell homing receptors (CCR10, CCL28 and α_4_β_7_-integrin) throughout mucosal surfaces, allowing circulating activated B and T cells to be attracted to multiple mucosal sites (67). Here we have shown that intranasal immunization is more effective at reducing bacterial CFU in the bladder of mice. Intramuscular and subcutaneous routes of vaccination may be less effective because homing receptors are not expressed on B cells that are activated in peripheral lymph nodes (68). However, it has been shown that subcutaneous immunization performed with two weeks between boosts can significantly reduce kidney colonization and generate serum IgG in Balb/c mice when formulated with MPLA-FimH/MrpH (69), alum-FyuA (18) and adjuvant-IroN (17).

When administered intranasally, the adjuvant dmLT significantly reduced colonization of UPEC in the bladder during UTI. Other adjuvants were unable to produce equivalent results, independent of administration route. The success of dmLT recapitulates previous findings in experiments testing antigenicity of outer membrane iron receptors that utilized cholera toxin conjugated to antigens (27, 28). This is likely related to the similarity between the two toxins. Both are A_1_B_5_ toxins whose B subunits bind to ganglioside GM1 and facilitate endocytosis of the A subunit into the cytosol (70–72). In addition, we found that dmLT alone reduced colonization in the kidneys, consistent with other vaccines where dmLT was used as adjuvant (52). The mechanism of immune modulation for dmLT has been demonstrated to be a strong IL-17 response and activation of Th17 cells (52, 73), which may contribute to its success given the critical role of Th17 cells and IL-17 signaling during the host response to bladder infection (4, 28, 74).

The four antigens used to determine the optimal route and adjuvant for a urinary tract vaccine have similar cellular functions. These proteins display similar β-barrel structures, are located in the Gram-negative bacterial outer membrane, and mediate the uptake of siderophores (FyuA, IutA, and IreA) or heme molecules (Hma) from the extracellular environment. However, despite these similarities, there are marked differences in the host response when individually formulated with a single adjuvant. IutA was the only antigen to significantly reduce bacterial colonization in all sites of infection assessed. In addition, intranasal immunization with IutA produced the most robust serum IgG response independent of adjuvant. This could be due to the high percentage of its amino acid sequence likely to represent MHC class II epitopes predicted by BepiPred-2.0, when compared to Hma, IreA, and FyuA. Furthermore, IutA has been shown to be highly expressed in the mouse model of ascending UTI, as assessed by RNA microarray, and in women presenting with symptoms of cystitis (32, 75, 76). Indeed, RNAseq data from 14 different strains, stabilized immediately following urine collection, clearly demonstrated that *iutA, hma, fyuA*, and *ireA*, are highly expressed *in vivo* in women with acute cystitis (77). Based on its efficacy and antigenicity, IutA should be considered for inclusion in future UTI vaccines, particularly those aimed at increasing the breadth of protection against UPEC strains and overcoming the functional redundancy of iron receptors by incorporation of multiple antigens.

Traditional vaccine development has been focused on production of antibodies and immunological memory. Our intranasally administered vaccine using the adjuvant dmLT significantly reduces *E. coli* colonization of the murine bladder when formulated with Hma or IutA; however, serum antigen-specific antibody levels did not correlate with CFU in the current study using purified LPS-free proteins. These results could suggest that administration of adjuvants via our tested routes did not elicit an effective antibody response, and that cellular immunity may be involved in protection. To that point, we previously demonstrated that intranasal immunization with bacterial siderophores (yersiniabactin and aerobactin), that bind to our iron-receptor antigens, provides protection by an unknown non-antibody-mediated mechanism (16). In addition, experiments vaccinating mice with cholera toxin conjugated to outer-membrane iron receptors generated antigen-specific antibody universally, independent of any observed reduction in CFU during murine ascending UTI (28). Together these data suggest that a strong IgG antibody response is not indicative of protection against infection. Further work is required to identify the immune factors contributing to protection, which will accelerate the development of an effective human UTI vaccine.

## Materials and Methods

### Ethics statement

#### Animal protocols

All animal protocols were approved by the Institutional Animal Care and Use Committee (IACUC) at the University of Michigan Medical School (PRO00009173), and in accordance with the Office of Laboratory Animal Welfare (OLAW), the United States Department of Agriculture (USDA), and the guidelines specified by the Association for Assessment and Accreditation of Laboratory Animal Care, International (AAALAC, Intl.). Mice were anesthetized with a weight-appropriate dose (0.1 mL for a mouse weighing 20 g) of ketamine/xylazine (80-120 mg/kg ketamine and 5-10 mg/kg xylazine) by intraperitoneal injection. Mice were euthanized by inhalant anesthetic overdose followed by vital organ removal.

#### Bacterial strains and culture conditions

*E. coli* CFT073 was isolated from the blood and urine of a hospitalized patient with acute pyelonephritis and urosepsis (78) and *E. coli* 536 was isolated from a patient with pyelonephritis (79). Transurethral infections with CFT073 or 536 were performed in mice immunized with vaccine formulations containing Hma, IreA, IutA, or FyuA. *E. coli* CFT073 does not encode a functional FyuA protein, thus, strain 536 was used to challenge mice that were immunized with FyuA. Strains were cultured in lysogeny broth (LB: 10 g/L tryptone, 5 g/L yeast extract, 0.5 g/L NaCl) at 37°C with aeration until saturation or on LB agar at 37°C.

#### Antigen purification and concentration

Commercially produced purified antigens were supplied by GenScript as follows. DNA sequences of *hma, ireA, iutA* from *E. coli* CFT073 and *fyuA* from *E. coli* 536 were codon-optimized and individually synthesized in-frame with a 6X histidine affinity tag and subcloned into *E. coli* expression vector pET-30a. Recombinant plasmids were transformed into *E. coli* BL21 star (DE3), cultured with shaking at 37°C in TB medium containing kanamycin, induced with IPTG, and harvested by centrifugation at 8,000 rpm. Cell pellets were lysed by sonication. Following centrifugation at 13,000 rpm, the precipitate was dissolved using 8M urea. Target protein was filter sterilized using a 0.22 µm membrane, quantified by the Bradford protein assay, and protein purity was determined by SDS-PAGE and Western Blot. Purified protein was obtained from inclusion bodies with purity varying from 85-90%. Proteins were certified as being endotoxin-free with levels < 100 EU/mg. Prior to immunization and subsequent boosts protein was concentrated with 10,000 NMWL centrifugal filter units (EMD Millipore).

#### Vaccine formulation and administration

Formulation of vaccines by mixing antigen and adjuvant was performed on the day of primary immunization and on the day of each subsequent boost. For each trial, seven to eight-week old CBA/J mice are given a primary dose of adjuvant alone, or adjuvant combined with 100 µg purified, LPS-free antigen via the specified route, intramuscular (IM), subcutaneous (SQ), or intranasal (IN), on day 0. On days 7 and 14, mice are boosted with adjuvant alone, or adjuvant combined with 25 µg antigen by the same route. On day 21 blood is collected retro orbitally via capillary tube, and mice are transurethrally inoculated with a UPEC strain expressing the antigen of interest. Forty-eight hours post inoculation urine is collected via abdominal massage and mice are sacrificed. Bladder and kidneys are harvested and bacterial burden per mL urine or per gram tissue determined. Amount and concentration of adjuvant for each vaccine was based upon pre-determined values as noted in product specification sheets and in published studies (80–83). A total of five adjuvants were tested for their efficacy within the vaccine formulations: Aluminum hydroxide gel (alum) (Alhydrogel adjuvant 2%, InvivoGen), Polyinosinic-polycytidylic acid (poly (I:C) HMW) (InvivoGen), Monophosphoryl Lipid A from *Salmonella minnesota* R595 (MPLA-SM) (InvivoGen), unmethylated CpG synthetic oligodeoxynucleotides ODN 2395, type C (CpG) (InvivoGen), and double mutant heat-labile toxin 1LT(R192G/L211A) (dmLT) (provided by Dr. John D Clements and Dr. Elizabeth Norton; Tulane University School of Medicine) (84). See Supplemental Table 2 for dose of adjuvant administered for each immunization route. For each vaccine trial, a control vaccine formulation was prepared containing adjuvant alone and phosphate buffered saline (PBS) with 1mM ethylenediamine tetraacetic acid (EDTA). Prepared antigen formulations were administered to seven to eight-week old female CBA/J mice IN (20 μL/mouse, 10 μL/nare), IM (50 μL/mouse) or SQ (70 μL/mouse).

#### Murine model of ascending UTI

Female CBA/J mice were transurethrally challenged as previously described (85). Briefly, bacterial pellets were harvested with centrifugation (3000 x *g*, 30 min, 4°C) and resuspended in phosphate-buffered saline (PBS: 8 g/L NaCl, 0.2 g/L KCl, 1.44 g/L Na_2_HPO_4_, 0.24 g/L KH_2_PO_4_, pH 7.4) to a final dose of 2 x 10^9^ CFU/mL. Actual inocula for each experimental trial are compared in Supplemental Figure 1. Each mouse was anesthetized and transurethrally infected with 50 μL of bacterial suspension using a Harvard Apparatus with a flow rate of 100 μL/min. Forty-eight hours post inoculation blood and urine were collected, mice were euthanized and bladder and kidneys were harvested. Urine and organ homogenates were diluted, plated on LB agar using an Autoplate 4000 spiral plater (Spiral Biotech) and enumerated using a QCount automated plate counter (Spiral Biotech) to determine the CFU/mL urine or CFU/g tissue.

#### Antibody quantification by ELISA

Quantification of antigen-specific antibody concentrations via indirect enzyme-linked immunosorbent assay (ELISA) was performed as previously described (27). Briefly, 5 μg/mL purified protein diluted in bicarbonate/carbonate buffer (3.03 g/L Na_2_CO_3_, 6.0 g/L NaHCO_3_) was coated in each well and incubated at 4°C overnight. Plates were washed with PBST (PBS containing 0.05% Tween 20) using an ELx405 microplate washer (Bio-Teck Instruments, Inc.) and blocked with SuperBlock (Pierce). Following a second wash in PBST, 50 μL of sera diluted in SuperBlock or undiluted urine were added to wells and incubated for 1-2 h at room temperature. Plates were again washed with PBST and coated with 50 μL 1:10,000 diluted secondary antibody goat anti-mouse IgG HRP conjugated (1030-05, SouthernBiotech) and incubated 1 hr at room temperature. After a final wash in PBST, 50 μL 1-Step Ultra TMB (3,3’,5,5’-tetramethylbenzidine) (Thermo Scientific) was added to each well and incubated at room temperature until sufficient color had developed. To stop the reaction, 50 μL 2M sulfuric acid was added to each well and the absorbance at 450 nm was read with a μQuant plate reader (Bio-Tek Instruments, Inc.). Antibody concentration was determined by comparing absorbance values to known concentrations of mouse IgG (0107-01, SouthernBiotech) bound to the plate with goat anti-mouse Ig (1010-01, SouthernBiotech). Serum assays were performed in duplicate for each mouse.

#### Statistical analysis

All graphic images and statistics were generated with Prism version 7 (GraphPad Software, Inc.). Significant differences in colonization levels and number of mice without detectable CFU were determined by a two tailed Mann-Whitney test, and Fisher’s exact test, respectively. Correlations between antibody concentrations and bacterial burden were determined by Pearson correlation coefficient.

## Acknowledgements

We would like to thank all members of the Mobley laboratory for their insightful comments and helpful critiques. We would also like to acknowledge Dr. John D. Clements and Dr. Elizabeth Norton for the gift of dmLT.

## Supplemental Figure Legends

**Supplemental Figure 1. Inocula are consistent across experimental trials and strains.** Inoculating doses of UPEC in CFU/mL administered transurethrally to female CBA/J mice one week following final boost. Mice were inoculated with strain CFT073 following immunization with Hma, IreA or IutA, or inoculated with strain 536 following immunization with FyuA. The intended dose was 2 x 10^9^ CFU/mL. Whiskers indicate maximum and minimum, box indicates 25^th^ and 75^th^ percentiles, bar indicates the median. N = 6 – 26. No statistical difference was found via two-tailed Mann-Whitney test.

**Supplemental Figure 2. Immunizing with CpG or polyIC in the absence of antigen does not affect colonization of the urinary tract by UPEC.** Seven to eight-week old female CBA/J mice were immunized intranasally according to our immunization schedule with the adjuvants: unmethylated CpG synthetic oligodeoxynucleotides (CpG), detoxified *E. coli* enterotoxin (dmLT), or polyinosinic:polycytodylic acid (polyIC). One week after the final boost, mice were transurethrally challenged with 10^8^ CFU of CFT073. Forty-eight hours post inoculation, urine was collected, mice were sacrificed, bladder and kidneys homogenized, and aliquots plated on LB agar for enumeration of bacterial burden. Bars indicate the median CFU in the urine (A), bladder (B), and kidneys (C). Symbols represent individual mice. N = 5-40. The limit of detection is 100 CFU/mL urine or /g tissue. Dotted line represents colonization level observed in unimmunized mice challenged with CFT073. *P* values shown where addition of adjuvant was protective as determined using a two-tailed Mann-Whitney test.

**Supplemental Figure 3. Subcutaneous vaccination with outer membrane iron receptors generates an antigen-specific serum IgG response.** Antigen-specific IgG concentrations quantified by indirect ELISA in pooled serum collected from female CBA/J mice one week after final boost. Mice were subcutaneously immunized with adjuvant alone (Ctl) or adjuvant formulated with 100 µg of purified, LPS free antigen (Hma, IreA, FyuA, IutA). Adjuvants utilized were: detoxified *E. coli* enterotoxin, (dmLT), polyinosinic:polycytodylic acid (polyIC), alum, and monophosphoryl lipid A (MPLA). Each bar represents the mean from four technical replicates of serum pooled from 10 mice from either one or two experimental trials, error bars indicate standard deviation. *P* values were determined using a two-tailed Mann-Whitney test. ** *P* ≤0.01, * *P* ≤0.05.

**Supplemental Figure 4. Antigen-specific serum antibodies do not correlate with protection against UTI.** The CFU/mL urine or /g tissue (y-axis) of all vaccination trials that significantly reduced bacterial burden (see Figure 1 and 3) were correlated with antigen-specific serum IgG concentration measured by ELISA (x-axis). Seven to eight-week old female CBA/J mice were immunized intranasally according to our immunization schedule with detoxified *E. coli* enterotoxin (dmLT) formulated with Hma (A), dmLT-IutA (B), polyinosinic:polycytodylic acid formulated with IutA (C), or unmethylated CpG synthetic oligodeoxynucleotides formulated with IutA (D). Each symbol represents a single mouse. Open symbols represent mice immunized with adjuvant alone, closed symbols represent mice immunized with adjuvant formulated with antigen. N = 5-17. No significant correlations were found using Pearsons correlation coefficient.

**Supplemental Table 1. Percent of immunized mice without detectable CFU following transurethral challenge.**

**Supplemental Table 2. Doses for each adjuvant by route of administration.**

